# Inter- and intra-specific variation of spider mite susceptibility to fungal infections: implications for the long-term success of biological control

**DOI:** 10.1101/615344

**Authors:** Flore Zélé, Mustafa Altıntaş, Inês Santos, Ibrahim Cakmak, Sara Magalhães

## Abstract

Spider mites are severe pests of several annual and perennial crops worldwide, often causing important economic damages. Moreover, rapid evolution of pesticide resistance in this group hampers the efficiency of chemical control. Alternative control strategies, such as the use of entomopathogenic fungi, are thus being developed. However, while several studies have focussed on the evaluation of the control potential of different fungal species and/or isolates as well as their compatibility with other control methods (e.g. predators or chemical pesticides), knowledge on the extent of inter- and intraspecific variation in spider mite susceptibility to fungal infection is as yet incipient. Here, we measured the mortality induced by two generalist fungi, *Beauveria bassiana* and *Metarhizium brunneum*, in 12 spider mite populations belonging to different *Tetranychus* species: *T. evansi*, *T. ludeni*, the green form of *T. urticae* and the red form of *T. urticae*, within a full factorial experiment. We found that spider mite species differed in their susceptibility to infection to both fungal species. Moreover, we also found important intraspecific variation for this trait. These results draw caution on the development of single strains as biocontrol agents. Indeed, the high level of intraspecific variation suggests that (a) the one-size-fits-all strategy will probably fail to control spider-mite populations and (b) hosts resistance to infection may evolve at a rapid pace. Finally, we propose future directions to better understand this system and improve the long-term success of spider mite control strategies based on entomopathogenic fungi.

## INTRODUCTION

Pesticides are still the main weapon used to control crop pests and disease vectors, despite the major threats they represent for food safety and for the environment (Bourguet and Guillemaud 2016). Moreover, the pervasive evolution and rapid spread of resistance to pesticides severely affect their efficiency in many taxa (Casida and Quistad 1998). Therefore, alternative control strategies are being sought to control disease epidemics and outbreaks of agricultural crop pests (Hajek et al. 2007; Lacey et al. 2001; Parolin et al. 2012; Zindel et al. 2011), including spider mites (Attia et al. 2013).

Spider mite of the genus *Tetranychus* (Acari: Tetranychidae) are ubiquitous major crop pests of c.a. 1100 plant species belonging to more than 140 different plant families (Migeon and Dorkeld 2006-2017), destroying annual and perennial crops. A few studies have evaluated the economic costs of spider mites, which vary among crops, seasons and plant age (Alatawi et al. 2007; Flamini 2006; Opit et al. 2005; Park and Lee 2002; Park and Lee 2005; Park and Lee 2007; Weihrauch 2005), and mathematical models suggest that detrimental effects of spider mites in agriculture will dramatically increase with increased global warming (Migeon et al. 2009). Moreover, due to their short generation time and high fecundity, spider mites rapidly develop resistance to pesticides (Van Leeuwen et al. 2010). Important efforts are thus being developed to evaluate the efficiency of different biological control methods, such as the use of essential oils, but also natural enemies such as predators, entomopathogenic bacteria and fungi (Attia et al. 2013). In particular, a plethora of studies have evaluated the virulence of many fungal species (e.g. *Neozygites* spp., *Metarhizium* spp., *Beauveria bassiana* and *Lecanicillium lecanii*) and/or strains to identify the best candidates for efficient spider mite control (e.g. Bugeme et al. 2008; Chandler et al. 2005; Maniania et al. 2008; Shi et al. 2008; Shin et al. 2017), as well as their compatibility with other control methods, such as predatory mites (e.g. Dogan et al. 2017; Seiedy 2014; Seiedy et al. 2012; Seiedy et al. 2013; Ullah and Lim 2017; Vergel et al. 2011; Wu et al. 2016) or pesticides (e.g. Irigaray et al. 2003; Klingen and Westrum 2007; Shi et al. 2005). However, these studies were conducted using a single host population and potential intraspecific variations in spider mites susceptibility have, to our knowledge, never been investigated within a single experiment (but see, for instance, Afifi et al. 2010; Fiedler and Sosnowska 2007; Ribeiro et al. 2009 for a comparison among spider mites and/or among other arthropod species; or Milner 1982; 1985; Perinotto et al. 2012; Uma Devi et al. 2008 for intraspecific variation within other arthropod species).

Both intra- and interspecific variability in host susceptibility to infection may modify epidemiological patterns of parasite in natural host populations (Dwyer et al. 1997; Hawley and Altizer 2011; Read 1995), thereby altering the efficiency and environmental persistence of biocontrol agents. Moreover, the use of such agents generates a strong selection pressure on the target pests (e.g. Fenner and Fantini 1999; see also Moscardi 1999; Tabashnik 1994) and, in general, variability in host susceptibility to infection may have important consequences for the evolution of host resistance as well as parasite virulence and transmission (Elena 2017; Sorci et al. 1997; Stevens and Rizzo 2008). Hence, assessing both intra- and interspecific variability in spider mite susceptibility to infection by different potential biocontrol agents is a prerequisite for the development of efficient and long-lasting control strategies.

Here, we assessed the susceptibility to fungi infection of 12 different spider mite populations belonging to different species that are ubiquitous in Europe and often co-occur in the field (Migeon and Dorkeld 2006-2017; Zélé et al. 2018b): 3 populations of the green form of *T. urticae*, 3 populations of the red form of *T. urticae* (also referred to as *T. cinnabarinus* by some authors; e.g. Li et al. 2009; Shi et al. 2005; Shi and Feng 2004), 3 populations of *T. ludeni*, and 3 populations of *T. evansi*. We used two generalist entomopathogenic fungi species, *Beauveria bassiana* and *Metarhizium brunneum*, as *Beauveria* and *Metarhizium* spp. are among the most used fungi in commercial production (Vega et al. 2009), and have wide geographical and host ranges (Greif and Currah 2007; Gurlek et al. 2018; Meyling and Eilenberg 2007; Rehner 2005; Roberts and Leger 2004). We then discuss the possible ecological and evolutionary causes and underlying mechanisms leading to the observed results, as well as their potential consequences for the evolution of both hosts susceptibility to infection and fungi virulence. Finally, we propose future directions to improve long-term success of spider mite control strategies using entomopathogenic fungi.

## MATERIALS AND METHODS

### Spider mite populations and rearing

Twelve populations of Tetranychid mites were used in this study. Three of *T. evansi* (called BR, GH and QL), three of *T. ludeni* (called Obi, Alval and Assaf), three of the red form of *T. urticae* (called AlRo, AMP.tet, FR.tet), and three of the green form of *T. urticae* (called TOM.rif, LS.tet, B6JS). Most of these populations were collected in Portugal from 2013 to 2016, FR.tet was collected in France and AlRo in Spain in 2013. The population BR of *T. evansi* was collected in a greenhouse in Brazil in 2002 (Godinho et al. 2016; Sarmento et al. 2011), and the population LS.tet of the green form of *T. urticae* derived from the London strain, which was used to sequence the species genome (Grbic et al. 2011). These populations originated from various plant species in the field, and none of them carried bacterial endosymbionts (i.e. *Wolbachia*, *Cardinium*, *Rickettsia*, *Arsenophonus*, *Spiroplasma*), either because they were initially uninfected when collected in the field (Zélé et al. 2018a), or following antibiotic treatment (3 generations with tetracycline hydrochloride, or 1 generation with rifampicin; all populations with “.tet” or “.rif” suffix, respectively; Breeuwer 1997; Gotoh et al. 2005; Li et al. 2014). All the information concerning these populations is summarized in Table S1. They were subsequently reared in the laboratory under standard conditions (25 ± 2°C, 60% RH, 16/8 h L/D) at high numbers (c.a. 500-1000 females per cage) in insect-proof cages containing either bean cv. Contender seedlings (obtained from Germisem, Oliveira do Hospital, Portugal) for *T. urticae* and *T. ludeni*, or tomato cv. Money Maker seedlings (obtained from Mr. Fothergill’s Seeds, Kentford, UK) for *T. evansi*.

### Entomopathogenic fungi strains and preparation of inoculum

We used the strains V275 (= Met52, F52, BIPESCO 5) of *Metarhizium brunneum* and UPH-1103 of *Beauveria bassiana*, obtained from Swansea University (UK) and from Siedlce University (Poland), respectively, as they were previously shown to have the potential to suppress *T. urticae* populations (Dogan et al. 2017). The procedures used for fungal growth, inoculum preparation and spider mite infection are similar to that described in Dogan et al. (2017). Briefly, the two fungi were grown on Sabouraud Dextrose Agar (SDA) medium at 25 °C for 2 weeks. Conidia were harvested from sporulating cultures with the aid of a spatula, washed with sterile distilled water and filtered through 4 layers of gauze to remove any hyphae.

### Spider mite infection and survival

The experiment was conducted in a growth chamber under standard conditions (25 ± 2°C, 80% RH, 16/8 h L/D). Roughly 2 weeks prior to the experiment, age cohorts were created for each spider mite population by collecting ca. 100 females from each mass culture, allowing them to lay eggs during 4 days on detached bean leaves placed on water-soaked cotton. The offspring from these cohorts was used in the experiment.

One day prior to the onset of this experiment, 20 adult mated females with similar age were randomly collected from each cohort of each population and placed on a 9-cm^2^ bean leaf disc placed on wet cotton (to ensure the leaf remained hydrated) with the abaxial (underside) surface facing upwards. On the first day of the experiment, the surface of the leaf discs was sprayed using a hand sprayer with 2.5 ml of a spore suspension of *M. brunneum* or *B. bassiana* in 0.03% (v/v) aqueous Tween 20 at 1 × 10^7^ conidia/ml, or, as control, with 0.3% aqueous Tween 20 only. Twelve replicates per treatment per population were performed within 7 temporal blocks (roughly 3 replicates of each treatment per block).

### Statistical analysis

Analyses were carried out using the R statistical package (version 3.5.3). Survival data were analysed using Cox proportional hazards mixed-effect models (coxme, kinship package). Spider-mite species, or populations within each species, and infection treatment (sprayed with *Beauveria bassiana*, with *Metarhizium brunneum*, or with Tween 20 only as control) were fit in as fixed explanatory variables, whereas discs nested within population, population (in the case of interspecific variation only) and block were fit as random explanatory variables. Hazard ratios (HR) were obtained from these models as an estimate of the difference between the rates of dying (i.e. the instantaneous rate of change in the log number of survivors per unit time; (Crawley 2007) between the controls of each species/population (by changing the intercept of the model) and the BB or MB treatments.

Maximal models, including all higher-order interactions, were simplified to establish a minimal model by sequentially eliminating non-significant terms and interactions (Crawley 2007). The significance of the explanatory variables was established using chi-squared tests (Bolker 2008). The significant chi-squared values given in the text are for the minimal model, whereas non-significant values correspond to those obtained before deletion of the variable from the model.

To further explore significant interactions between species/population and treatment effects on HR, the two factors were concatenated to fit a single fixed factor containing all species/population by treatments levels in the models. Multiple comparisons were then performed using General Linear Hypotheses (glht, package multicomp) with Holm corrections.

## RESULTS

### *Interspecific variation of spider-mite susceptibility to infection by* Beauveria bassiana *and* Metarhizium brunneum

The statistical analyses revealed a significant interaction between treatments (females sprayed with either Tween 20 only as control, *B. bassiana*, or *M. brunneum*) and species (*T. evansi*, *T. ludeni*, red and green form of *T. urticae*) on the survival of spider mites (*X*^*2*^_*6*_=80.61, p<0.001; Fig.1 a-d). Multiple comparisons of hazard ratios (HRs) obtained for each spider-mite species infected by each fungal species relative to the control revealed that all species were not equally affected by infection (Fig. 1e): both fungi induced a stronger mortality in *T. evansi* (HR=5.04 for *B. bassiana*, and HR=5.15 for *M. brunneum*) and in the green form of *T. urticae* (HR=5.30 for *B. bassiana*, and HR=6.27 for *M. brunneum*), than in *T. ludeni* (HR=3.25 for *B. bassiana*, and HR=3.73 for *M. brunneum*) and in the red form of *T. urticae* (HR=3.84 for *B. bassiana*, and HR=3.21 for *M. brunneum*); but their effect did not differ within these two groups of species (Fig. 1e; cf. Table S1 for the statistical results of all multiple comparisons). Moreover, while the two fungi induced similar levels of mortality in *T. evansi* (z=−0.39, p=1.00) and in *T. ludeni* (z=−0.14, p=0.13), infection with *B. bassiana* led to higher mortality than that with *M. brunneum* in the red form of *T. urticae* (z=0.18, p=0.03), while the reverse was found in the green form of *T. urticae* (z=−0.17, p=0.04). Note that survival in the *T. evansi* control was higher than in that of the three other species (Fig. S1 and Table S2).

**Figure 1.**
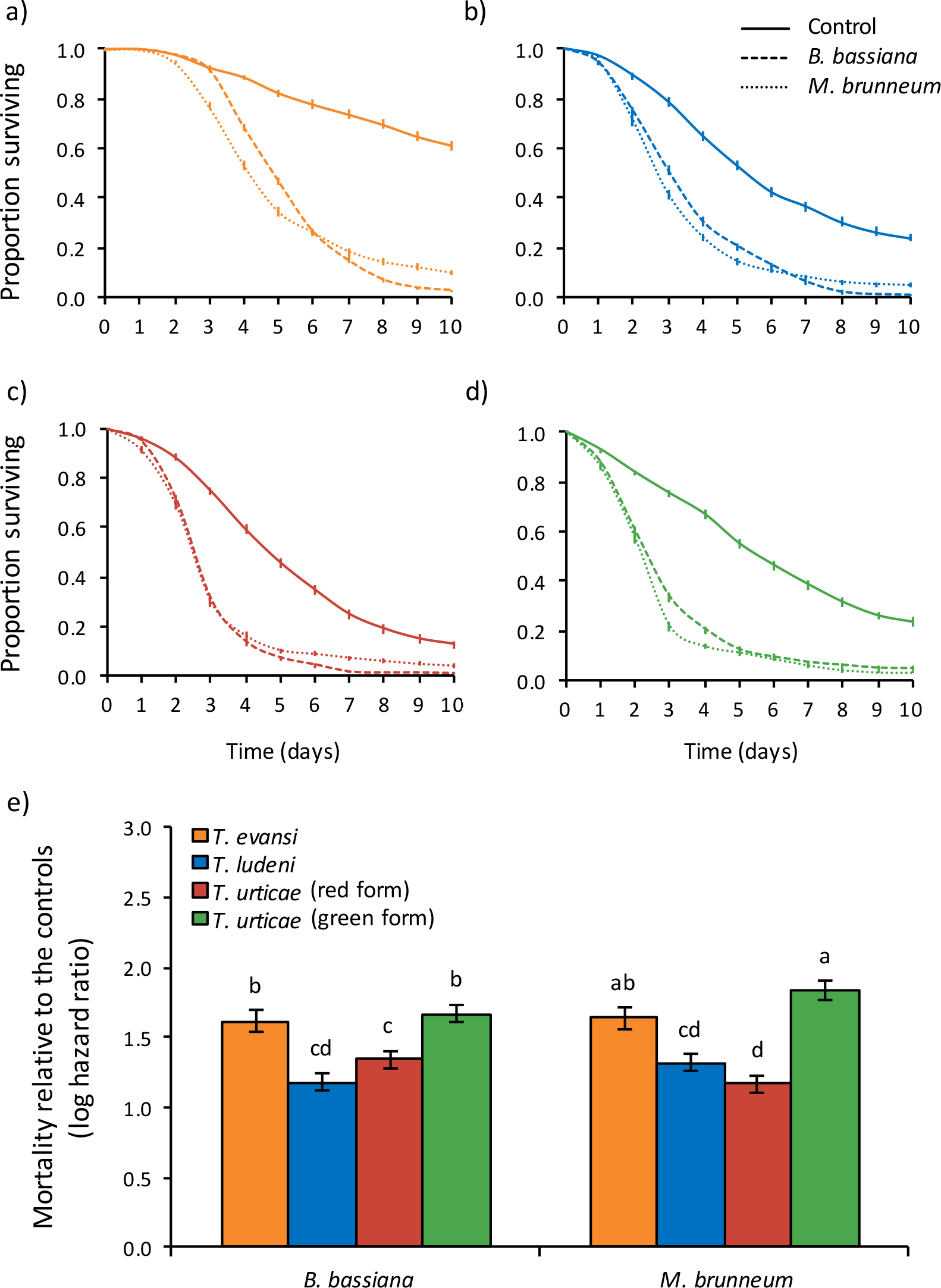
Survival curves of spider-mite females from (a) *T. evansi,* (b) *T. ludeni,* (c) red form of *T. urticae,* and (d) green form of *T. urticae*, sprayed with *B. bassiana* (dashed lines), *M. brunneum* (dotted lines), or with Tween 20 only (control; solid lines). (e) mortality upon infection by each fungus relative to the controls (log hazard ratio ± s.e.); identical letter superscripts indicate non-significant differences between treatments at the 5% level (multiple comparisons with Holm correction).

### *Intraspecific variation of spider-mite susceptibility to infection by* Beauveria bassiana *and* Metarhizium brunneum

We also found a significant interaction between treatment and population on spider-mite survival within each of the species studied (in the green form of *T. urticae*: *X*^*2*^_*4*_=79.60, p<0.0001; in the red form *T. urticae*: *X*^*2*^_*4*_=12.12, p<0.02; in *T. ludeni*: *X*^*2*^_*4*_=17.41, p<0.002; in *T. evansi*: *X*^*2*^_*4*_=106.72, p<0.0001). Indeed, for all species, the spider mite populations differed in their susceptibility either to both fungi or to only one of them.

In the green form of *T. urticae*, *B. bassiana* induced a higher mortality in the populations TOM.rif and B6JS (a 6.01-fold and a 4.41-fold decrease in survival upon infection relative to the control, respectively) than in the population LS.tet (a 2.85-fold decrease in survival). Similarly, *M. brunneum* induced higher mortality in B6JS (a 8.67-fold decrease in survival) than in TOM.rif (a 4.60-fold decrease in survival), and the highest mortality was found in LS.tet (a 3.04-fold decrease in survival; Fig. 2a-d; see Table S3 for the statistical results of all comparisons). Moreover, while infection with both fungi led to similar mortality in LS.tet (z=−0.67, p=1.00), infection with *B. bassiana* induced a higher mortality rate than that with *M. brunneum* in TOM.rif (z=2.87, p=0.04), and the reverse was found in B6JS (z=−6.86, p<0.0001).

**Figure 2.**
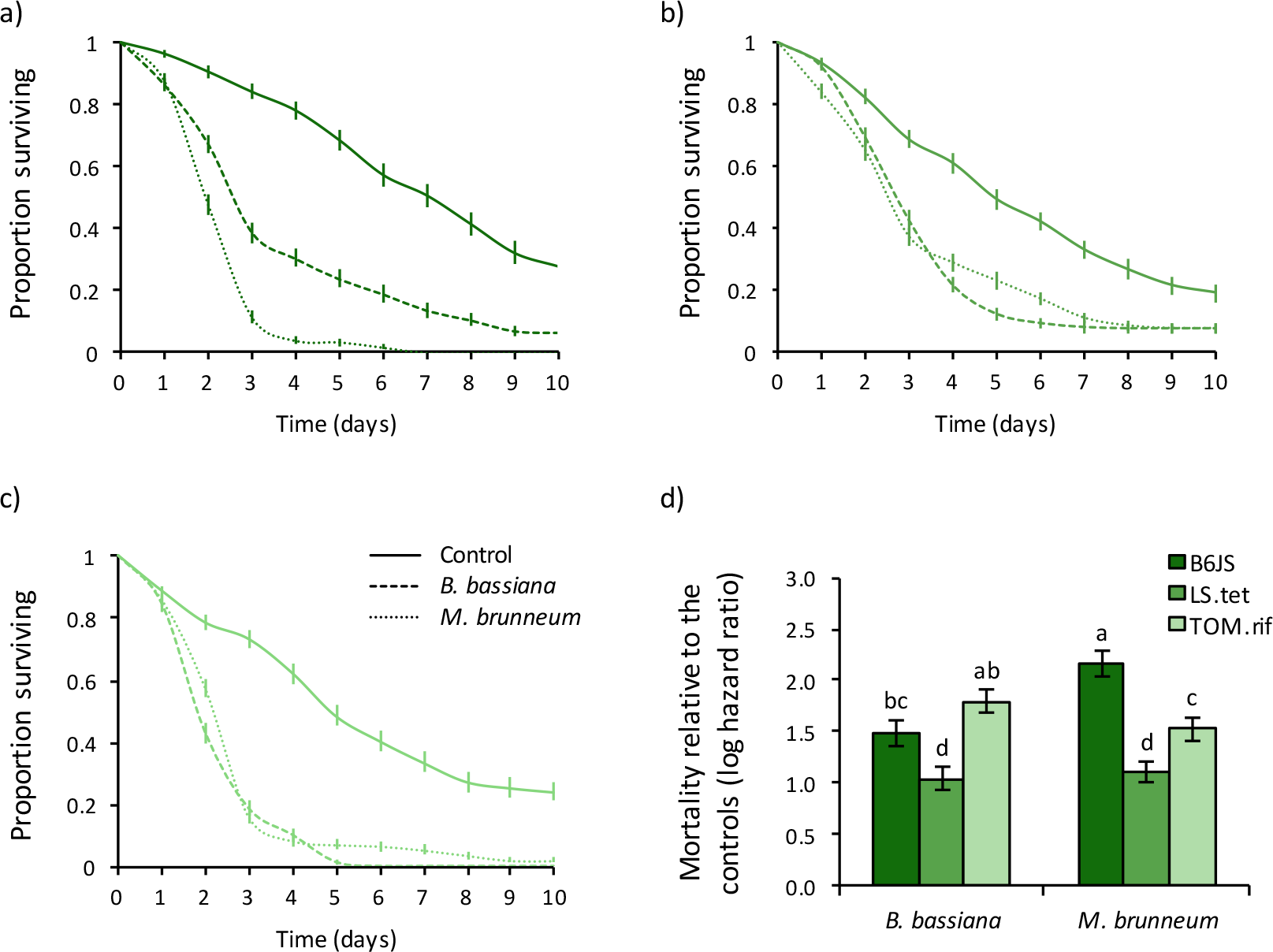
Survival curves of spider-mite females from different populations of the green form of *T. urticae*: (a) B6JS, (b) LS.tet, and (c) TOM.rif, sprayed with *B. bassiana* (dashed lines), *M. brunneum* (dotted lines), or with Tween 20 only (control; solid lines). (d) mortality relative to the controls (log hazard ratio ± s.e.); identical letter superscripts indicate non-significant differences between treatments at the 5% level (multiple comparisons with Holm correction).

**Figure 3.**
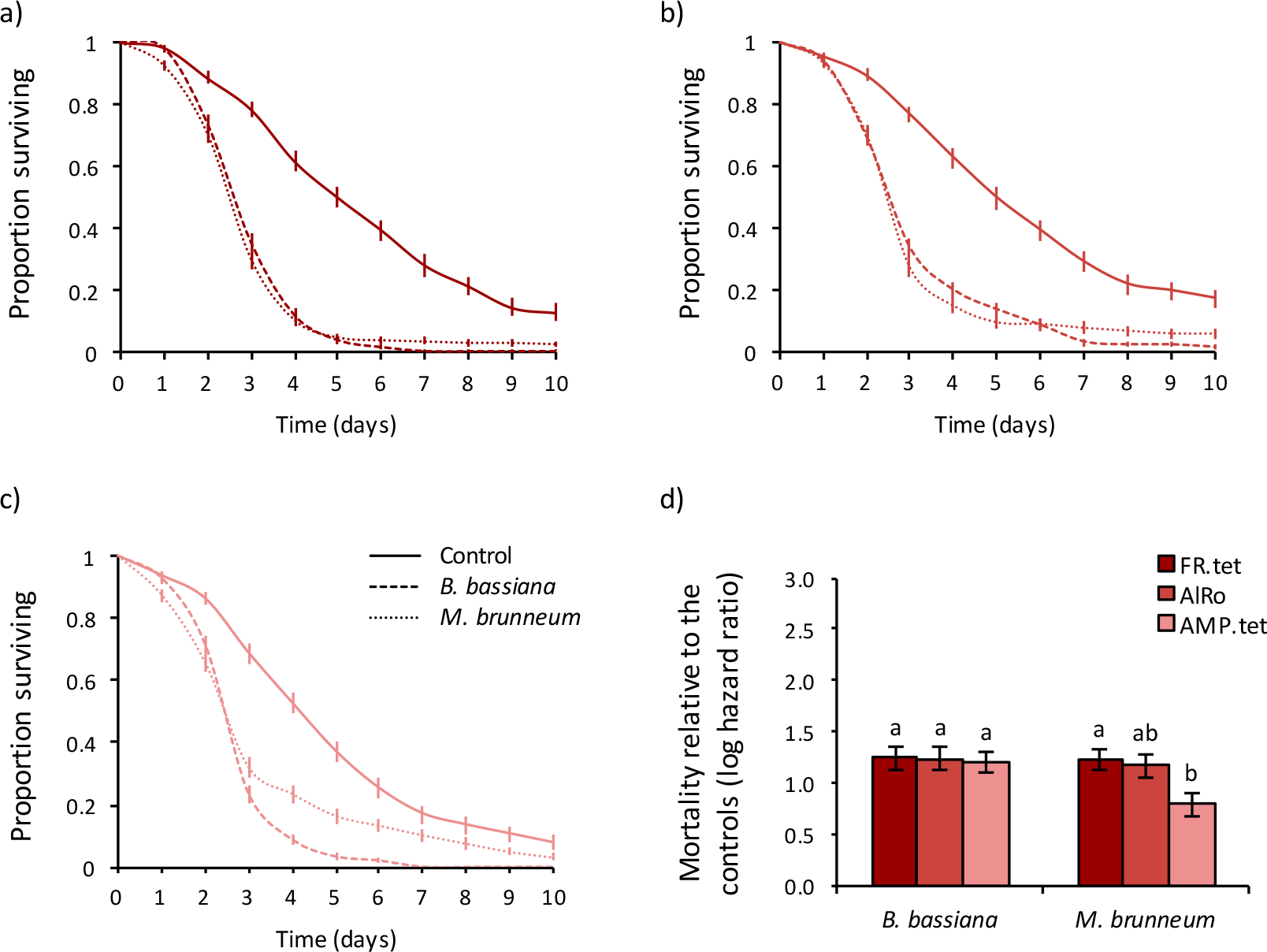
Survival curves of spider-mite females from different populations of the red form of *T. urticae*: (a) FR.tet, (b) AlRo, and (c) AMP.tet, sprayed with *B. bassiana* (dashed lines), *M. brunneum* (dotted lines), or with Tween 20 only (control; solid lines). (d) mortality relative to the controls (log hazard ratio ± s.e.); identical letter superscripts indicate non-significant differences between treatments at the 5% level (multiple comparisons with Holm correction).

In the red form of *T. urticae*, both fungi had the same effect in all populations (HR ranged between 3.21 and 3.47) except in the population AMP.tet in which *M. brunneum* induced lower mortality (HR=2.21; Fig. 3a-d; see Table S4 for the statistical results of all comparisons).

In *T. ludeni*, both fungi induced similar mortality in all populations except for the population Alval, in which *M. brunneum* induced higher mortality than in the population Assaf, with a 4.86-fold and 3.17-fold decreased survival upon infection relative to the control, respectively (HR ranged between 2.98 and 3.77 in the other treatments; Fig. 4a-d; see Table S7 for the statistical results of all comparisons). Note that, in this species, the population controls (not exposed to fungi) did not have the same survival (Fig. S1; Table S5).

In *T. evansi*, both fungi species induced similar mortality independently of the population tested, but they this induced mortality was higher in the populations GH and BR than in the population QL (c.a. 12-fold, 10-fold, and 2-fold decreased survival upon infection relative to the controls, respectively; Fig. 5a-d; see Table S6 for the statistical results of all comparisons). Note, however, that QL control had a much lower survival than that of the two other populations (Fig. S1; Tables S6).

**Figure 4.**
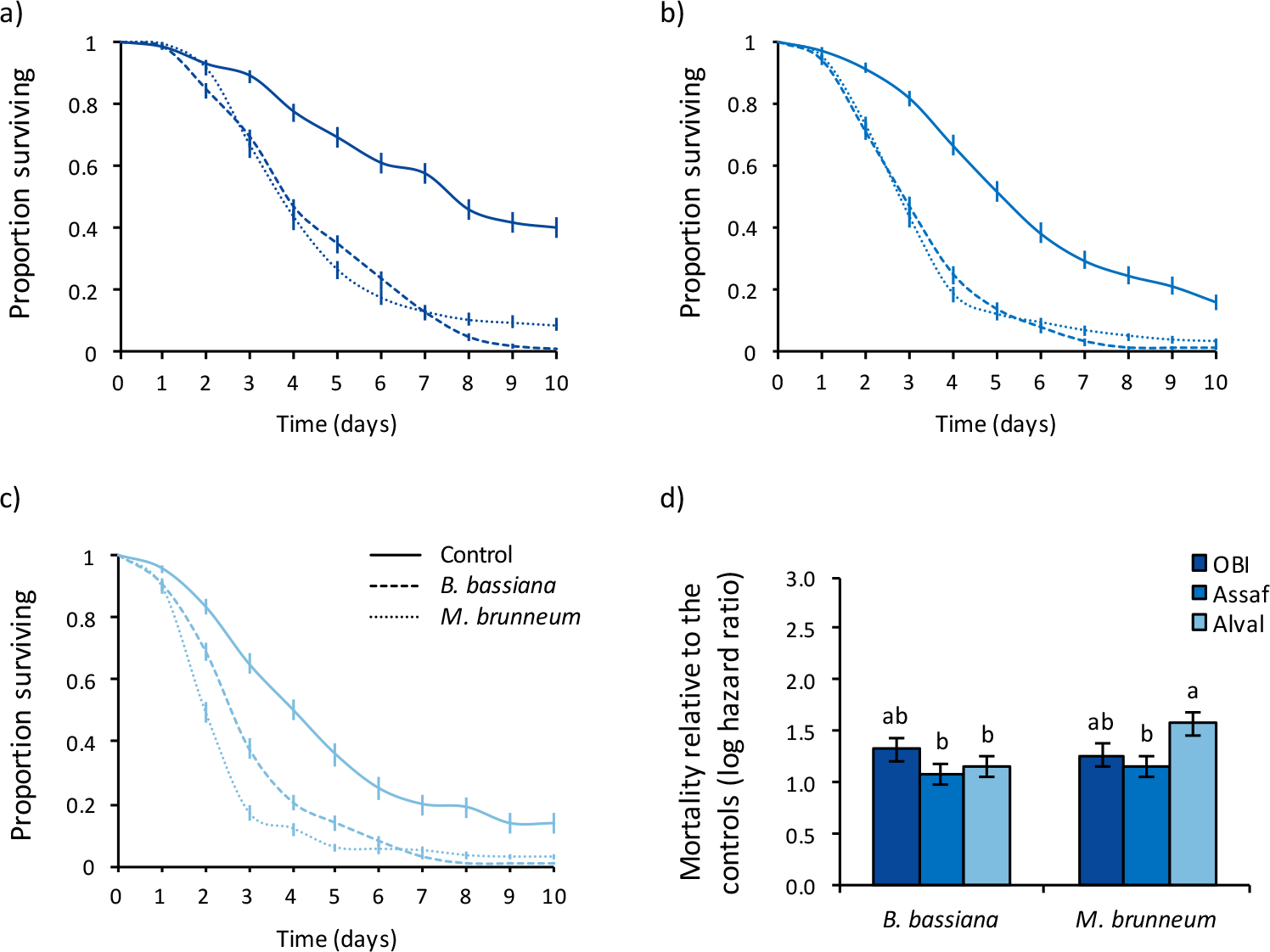
Survival curves of spider-mite females from different *T. ludeni* populations: (a) OBI, (b) Assaf, and (c) Alval, sprayed with *B. bassiana* (dashed lines), *M. brunneum* (dotted lines), or with Tween 20 only (control; solid lines). (d) mortality relative to the controls (log hazard ratio ± s.e.); identical letter superscripts indicate non-significant differences between treatments at the 5% level (multiple comparisons with Holm correction).

**Figure 5.**
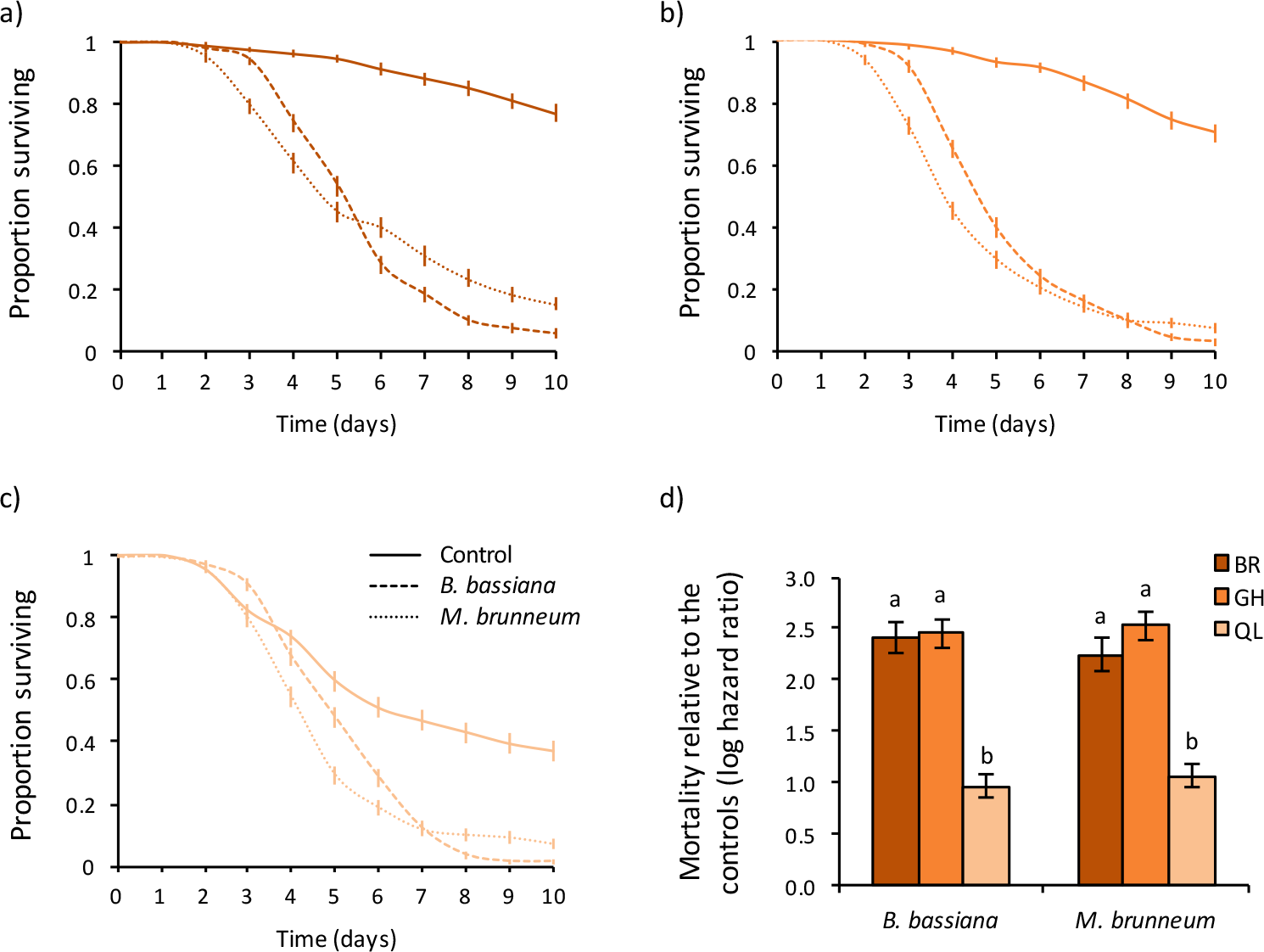
Survival curves of spider-mite females from different *T. evansi* populations: (a) BR, (b) GH, and (c) QL, sprayed with *B. bassiana* (dashed lines), *M. brunneum* (dotted lines), or with Tween 20 only (control; solid lines). (d) mortality relative to the controls (log hazard ratio ± s.e.); identical letter superscripts indicate non-significant differences between treatments at the 5% level (multiple comparisons with Holm correction).

## DISCUSSION

In this study, we found both intra- and interspecific variability in the susceptibility of *Tetranychus* spider mite to infection by *B. bassiana* and *M. brunneum*. Overall, we observed a higher mortality upon infection in *T. evansi* and in the green form of *T. urticae*, than in *T. ludeni* and in the red form of *T. urticae*. These results, however, may not reflect accurately the virulence of both fungi in each of these spider mite species. Indeed, we further found important variation among populations within each species. Most variation was found among populations of *T. evansi* and of the green form of *T. urticae*, with, for instance, the mortality upon infection of two populations of *T. evansi* (BR and GH) being 5 times higher than that of another (QL). We also found variation among populations of *T. ludeni* and of the red form of *T. urticae*, although the amplitude of these effects was relatively smaller and depended on the fungal species.

Overall, our results suggest that spider mite susceptibility to infection is not a phylogenetically-conserved trait, and further corroborate the generalist status of both fungal species (Meyling and Eilenberg 2007; Rehner 2005; Roberts and Leger 2004). For instance, *B. bassiana* occurs naturally in more than 700 host species (Inglis et al. 2001), and this range is likely underestimated as prevalence estimates are usually done in arthropod species that are crop pests or predators and parasitoids used as biocontrol agents (Meyling and Eilenberg 2007). Moreover, differences in virulence between the two fungi shown here suggest population-specific responses to each fungus, instead of a more general response against infection. For instance, *M. brunneum* is more virulent than *B. bassiana* in the population B6JS of green *T. urticae* and in the population Alval of *T. ludeni*, while the reverse occurred in the population AMP.tet of the red form of *T. urticae*. Such differences in susceptibility to infection between populations independently of their phylogenetic relationship may thus reflect differences in exposure by each fungus species (i.e. different selection pressure for resistance mechanisms to evolve) throughout their evolutionary history.

Variations in the prevalence of each fungus leading to different exposure may, for instance, occur between different geographical areas due to several environmental factors, such as temperature, humidity and solar (UV) irradiation (Meyling and Eilenberg 2007). However, these fungi are known to have a cosmopolitan distribution and our results show no clear association between the susceptibility of a particular spider mite population and its country of origin. For instance, the *T. evansi* populations BR and GH come from Brazil and Portugal, respectively, but do not differ in susceptibility to infection by both fungi; similarly, the effect of *B. bassiana* do not differ between populations of red *T. urticae* collected in France (FR.tet), Spain (AlRo) and Portugal (AMP.tet). Instead, we found different susceptibility to infection between populations at small geographical scales, such as in the *T. evansi* populations GH and QL and in the green *T. urticae* populations B6JS and TOM.rif upon infection by both fungi; or in the *T. ludeni* populations Assaf and Alval upon infection by *M. brunneum*; while all of these populations were collected in the same region in Portugal. These results might thus be explained by microhabitats-specific distribution of the fungi, as previously found for different isolates of *B. bassiana* (e.g. Ormond et al. 2010; Wang et al. 2003). Moreover, several studies suggest that both *B. bassiana* and *Metarhizium* spp. have the potential to interact directly with the host plants of arthropods (reviewed in Meyling and Eilenberg 2007), which may potentially lead to plant-specific distribution of the fungi. Indeed, *Metarhizium* spp. occur in the rhizosphere, which possibly provides a “refuge” where the fungus can survive outside insect hosts, and the presence of *B. bassiana* in internal plant tissue has been discussed as an adaptive protection against herbivorous insects (reviewed in Meyling and Eilenberg 2007). However, the host plant range of these fungi is, to our knowledge, as yet unknown. Moreover, no field survey of these fungi has been conducted to date in *Tetranychus* spp. (but see, for instance, Debnath and Sreerama Kumar 2017; Dick and Buschman 1995; Van Der Geest et al. 2002, for other fungi and/or spider mite species). Future evaluation of the prevalence of infection by *M. brunneum* and *B. bassiana* in natural populations of spider mites collected on different host plants would thus be necessary to further understand possible factors that could explain the patterns observed in our experiment (Boots et al. 2009).

Decreased host susceptibility to infection may be the result of two different (albeit nonexclusive) mechanisms (Boots et al. 2009; Read et al. 2008): resistance (i.e. reduction in parasite load) and/or tolerance (i.e. reduction of the damage incurred by a parasite). Differential host resistance to fungal infection might be due, for instance, to variability in different cuticular barriers. Such barriers include the absence of factors necessary for parasite recognition, or the presence of inhibitory compounds (phenols, quinones, and lipids) on the cuticle surface, but also the cuticle thickness, its degree of hardening by sclerotization, its resistance to enzymatic degradation and its permeability (reviewed in Hajek and St. Leger 1994). Subsequently, when a fungus bypass cuticular barriers, variability in systemic immunity may also lead to differential host resistance responses. This may include differential activation of the Toll and JAK/STAT pathways, which converge into the transcriptional activation of genes involved in phagocytosis, encapsulation and humoral responses (e.g. Dong et al. 2012). Interestingly, several, but not all, important genes described in these pathways in *Drosophila melanosgaster* were absent in the genome of the green form of *T. urticae* (Grbic et al. 2011), and spider mites have high mortality upon bacterial infection (Santos-Matos et al. 2017). Whether the presence of such immune genes vary between or within spider mite species and their expression depend on different fungal species has not been explored to date. In particular, the absence of many important immune genes in *T. urticae* suggests that tolerance mechanisms (e.g. via a decrease of the immune response to avoid autophagy) rather than resistance have been favoured throughout their evolutionary history. However, such hypothesis remains to be tested and further studies are necessary to better understand the mechanisms of spider mite resistance and tolerance against fungal infection.

Independently of the underlying mechanisms at play, whether spider mite populations differ in resistance or tolerance to fungal infection may have different epidemiological and evolutionary consequences, and, hence, different implications for the long-term success of spider mite control. One the one hand, resistance to infection might be rapidly selected following application of fungi to crops, and subsequently invade spider mite populations, thereby decreasing fungi prevalence and hampering the success of such control strategy. On the other hand, host tolerance should have neutral or even positive effect on parasite prevalence (Boots et al. 2009; Miller et al. 2006; Read et al. 2008), but as, by definition, tolerance minimizes the harm caused by pathogens, it may hamper the efficiency of fungi in controlling spider mites. Moreover, host resistance and tolerance may lead to different evolutionary outcomes for parasite virulence (Boots et al. 2009). Indeed, whereas host resistance is predicted to select for increased parasite virulence (e.g.(Gandon and Michalakis 2000), host tolerance does not reduce parasite fitness and, therefore, will not lead to antagonistic counter-adaptation by pathogens (Raberg et al. 2007; Rausher 2001). Still, depending on the nature of the tolerance mechanism, it may lead to the evolution of more virulent and transmissible parasites (Miller et al. 2006), with potentially serious implications for non-tolerant populations (Boots et al. 2009), including non-target species such as crop auxiliaries or spider mite predators. Finally, although increased mortality due to infection should lead to a reduction in oviposition duration, spider mites may evolve the ability to compensate infection-driven fitness costs by changing the timing of their reproductive efforts (i.e. ‘fecundity compensation’; (Parker et al. 2011; Vezilier et al. 2015), thereby limiting the efficiency of fungi applications for population control. Hence, assessing which of these evolutionary outcome is more likely is timely. In particular, it is likely that the, the high level of intraspecific variation in susceptibility to infection found in our study is recapitulated within populations and is, at least partly, genetically determined. If this is the case, then this trait may evolve at a rapid pace.

In conclusion, our results show both intra- and interspecific variability in spider mite susceptibility to fungi-induced mortality using two generalist fungi, *B. bassiana* and *M. brunneum*. To our knowledge, this is the first study investigating the effect of entomopathogenic fungi on the survival of multiple spider mite populations belonging to different species within a single full factorial experiment. In line with laboratory virulence tests that are not necessarily well correlated with field effectiveness (Roberts and Leger 2004), our results highlight the importance of studying several host populations/genomes when assessing the efficiency of a given biocontrol agent, and draw caution on the development of single strains as biocontrol agents as hosts resistance to infection may evolve at a rapid pace.

## Supporting information

Electronic supplementary materials

## AUTHORS’ CONTRIBUTIONS

Experimental conception and design: FZ, SM; maintenance of spider mite populations and plants: IS; acquisition of data: MA; statistical analyses: FZ; paper writing: FZ, SM, with input from all authors. Funding: IC, SM. All authors have read and approved the final version of the manuscript.

## ACKNOWLEDGMENTS

We thank Diogo Godinho and Miguel Cruz for their help in some parts of the experiment, as well as Marta Palma for technical support. We also thank all members of the SM lab for useful discussions and suggestions. This work was funded by an FCT-Tubitak agreement (FCT-TUBITAK/0001/2014 and TUBITAK TOVAG 115O610) to IC and SM, and by Adnan Menderes University Research Foundation (ZRF-17055) to IC. FZ was funded through an FCT Post-Doc fellowship (SFRH/BPD/125020/2016). Funding agencies did not participate in the design or analysis of experiments. We declare that we do not have any conflict of interest.

