## Supplementary material for "Inter- and intra-specific variation of spider mite susceptibility to fungal infections: implications for the long-term success of biological control": Electronic supplementary materials

**Table S1.** Populations of spider mites used in the experiment. Mites were collected in Canada (C), Brazil (B), Portugal (P), France (F) and Spain (S). Populations harbouring bacterial endosymbionts (e.g. *Wolbachia*, *Cardinium*, *Rickettsia*) were cured from infection with antibiotics.

| Species | Name | Date | Host plant | Location | Coordinates | Antibiotics | Ref |
|---|---|---|---|---|---|---|---|
| *T. evansi* | BR | --/--/2002 | *Solanum lycopersicum* | Unknown (B) | unknown | - | [1] |
| | GH | 03/10/2013 | *Solanum lycopersicum* | University of Lisbon (P) | 38.757852, -9.158221 | - | [2] |
| | QL | 26/11/2013 | *Datura stramonium* | Quinta das Lameiras (P) | 39.085837, -8.991478 | - | [2] |
| *T. ludeni* | Obi | 09/10/2016 | *Ipomoea purpurea* | Óbidos, Gracieira (P) | 39.334049,-9.121178 | - | - |
| | Alval | 05/11/2013 | *Ipomoea purpurea* | Alvalade, Lisbon (P) | 38.75515, -9.14685 | - | [2] |
| | Assaf | 20/09/2013 | *Datura stramonium* | Assafora (P) | 38.904743, -9.408592 | - | [2] |
| *T. urticae* red form | AlRo | 09/11/2013 | *Rosa spp.* | Almería (S) | 36.855725, -2.320374 | - | [2] |
| | AMP.tet | 18/11/2013 | *Datura stramonium* | Aldeia da Mata Pequena (P) | 38.534363, -9.191163 | 13/01/2013[1] | [2] |
| | FR.tet | 11/10/2013 | *Solanum lycopersicum* | Montpellier (F) | 43.614951, 3.859846 | 26/12/2013[1] | [2] |
| *T. urticae* green form | TOM.rif | --/05/2010 | *Solanum lycopersicum* | Carregado (P) | 39.078962,-8.993656 | 15/09/2016[2] | [3] |
| | LS.tet | unknown | *Phaseolus vulgaris* | Vineland, Ontario (C) | unknown | 13/01/2013[1] | [4] |
| | B6JS[3] | 10/06/2015 | *Phaseolus vulgaris* | Correias (P) | 39.342914, -8.797936 | - | [5] |

[1] Tetracycline hydrochloride [6]
[2] Rifampicin [7]
[3] Population called B6 in the supplementary materials of [5]

**Table S2.** Results of multiple comparisons (with Holm correction) between hazard ratios obtained for different spider-mite species (Te: *T. evansi*; Tl: *T. ludeni*; TuR: red form of *T. urticae*; TuG: green form of *T. urticae*) among the different treatments (BB: *Beauveria bassiana*; MB: *Metarhizium brunneum*; C: Control). Hazard ratios of infection by each fungus were estimated relative to the control within each species.

| | Factor 1 | Factor 2 | Estimate | Std. Error | z value | Pr(>\|z\|) |
|---|---|---|---|---|---|---|
| Controls | Te_C | TuG_C | -0.710 | 0.204 | -3.472 | 0.008** |
| | TuG_C | Tl_C | -0.288 | 0.202 | -1.427 | 0.768 |
| | Tl_C | TuR_C | -0.303 | 0.196 | -1.541 | 0.740 |
| | TuG_C | TuR_C | -0.590 | 0.201 | -2.934 | 0.040* |
| BB | TuG_BB | Te_BB | 0.051 | 0.097 | 0.523 | 1.000 |
| | Tc_BB | Te_BB | -2.272 | 0.095 | -2.866 | 0.046* |
| | Tc_BB | Tl_BB | 0.166 | 0.086 | 1.927 | 0.432 |
| MB | Tu_MB | Te_MB | 0.196 | 0.097 | 2.011 | 0.399 |
| | Tl_MB | Te_MB | -0.323 | 0.097 | -3.337 | 0.013* |
| | Tc_MB | Tl_MB | -0.149 | 0.088 | -1.695 | 0.630 |
| BB vs MB | Te_BB | Te_MB | -0.022 | 0.056 | -0.391 | 1.000 |
| | Tl_BB | Tl_MB | -0.137 | 0.055 | -2.481 | 0.131 |
| | Tc_BB | Tc_MB | 0.179 | 0.057 | 3.128 | 0.025* |
| | Tu_BB | Tu_MB | -0.167 | 0.056 | -2.961 | 0.040* |
| | Tu_BB | Te_MB | 0.029 | 0.097 | 0.296 | 1.000 |
| | Tl_BB | Tc_MB | 0.013 | 0.087 | 0.148 | 1.000 |

**Table S3.** Results of the multiple comparisons (with Holm correction) between hazard ratios obtained for different populations of the green form of *T. urticae* (TOM.rif, LS.tet, and B6JS) among the different treatments (BB: *Beauveria bassiana*; MB: *Metarhizium brunneum*; C: control). Hazard ratios of infection by each fungus were estimated relative to the control within each population.

| | Factor 1 | Factor 2 | Estimate | Std. Error | z value | Pr(>\|z\|) |
|---|---|---|---|---|---|---|
| Controls | TOM.rif_C | LS.tet_C | -0.002 | 0.127 | -0.018 | 1.000 |
| | LS.tet_C | B6JS_C | 0.307 | 0.165 | 1.863 | 0.312 |
| | B6JS_C | TOM.rif_C | -0.305 | 0.169 | -1.809 | 0.312 |
| BB | B6JS_BB | TOM.rif_BB | -0.302 | 0.156 | -1.981 | 0.285 |
| | B6JS_BB | LS.tet_BB | 0.436 | 0.155 | 2.813 | 0.044* |
| MB | B6JS_MB | TOM.rif_MB | 0.634 | 0.154 | 4.118 | <0.0005*** |
| | TOM.rif_MB | LS.tet_MB | 0.414 | 0.147 | 2.816 | 0.044* |
| BB vs MB | TOM.rif_BB | TOM.rif_MB | 0.268 | 0.093 | 2.868 | 0.041* |
| | LS.tet_BB | LS.tet_MB | -0.065 | 0.096 | -0.671 | 1.000 |
| | B6JS_BB | B6JS_MB | -0.676 | 0.099 | -6.857 | 8.42e-11*** |
| | B6JS_BB | TOM.rif_MB | -0.042 | 0.156 | -0.271 | 1.000 |
| | TOM.rif_BB | B6JS_MB | -0.366 | 0.153 | -3.387 | 0.119 |

**Table S4.** Results of the multiple comparisons (with Holm correction) between hazard ratios obtained for different populations of the red form of *T. urticae* (AlRo, AMP.tet, and FR.tet) among the different treatments (BB: *Beauveria bassiana*; MB: *Metarhizium brunneum*; C: control). Hazard ratios of infection by each fungus were estimated relative to the control within each population.

| | Factor 1 | Factor 2 | Estimate | Std. Error | z value | Pr(>\|z\|) |
|---|---|---|---|---|---|---|
| Controls | AlRo_C | AMP.tet_C | -0.244 | 0.151 | -1.621 | 0.841 |
| | AMP.tet_C | FR.tet_C | 0.309 | 0.162 | 1.903 | 0.513 |
| | FR.tet_C | AlRo_C | -0.065 | 0.161 | -0.404 | 1.000 |
| BB | AlRo_BB | FR.tet_BB | -0.011 | 0.149 | -0.072 | 1.000 |
| | AlRo_BB | AMP.tet_BB | 0.037 | 0.147 | 0.248 | 1.000 |
| | FR.tet_BB | AMP.tet_BB | 0.047 | 0.146 | 0.324 | 1.000 |
| MB | AlRo_MB | FR.tet_MB | -0.062 | 0.148 | -0.416 | 1.000 |
| | AlRo_MB | AMP.tet_MB | 0.375 | 0.154 | 2.431 | 0.151 |
| | FR.tet_MB | AMP.tet_MB | 0.437 | 0.154 | 2.843 | 0.049* |
| BB vs MB | AlRo_BB | AlRo_MB | 0.068 | 0.098 | 0.690 | 1.000 |
| | AMP.tet_BB | AMP.tet_MB | 0.407 | 0.102 | 3.979 | 0.0008*** |
| | FR.tet_BB | FR.tet_MB | 0.017 | 0.094 | 0.182 | 1.000 |

**Table S5.** Results of the multiple comparisons (with Holm correction) between hazard ratios obtained for different *T. ludeni* populations (OBI, Assaf, and Alval) among the different treatments (BB: *Beauveria bassiana*; MB: *Metarhizium brunneum*; C: control). Hazard ratios of infection by each fungus were estimated relative to the control within each population.

| | Factor 1 | Factor 2 | Estimate | Std. Error | z value | Pr(>\|z\|) |
|---|---|---|---|---|---|---|
| Controls | OBI_C | Assaf_C | -0.639 | 0.156 | -4.088 | 0.0005*** |
| | Assaf_C | Alval_C | -0.141 | 0.144 | -0.980 | 1.000 |
| | Alval_C | OBI_C | 0.780 | 0.158 | 4.945 | 9.91e-06*** |
| BB | OBI_BB | Assaf_BB | 0.237 | 0.150 | 1.581 | 0.867 |
| | Assaf_BB | Alval_BB | 0.171 | 0.155 | 1.101 | 1.000 |
| | Alval_BB | OBI_BB | 0.066 | 0.147 | 0.447 | 1.000 |
| MB | OBI_MB | Assaf_MB | 0.113 | 0.153 | 0.738 | 1.000 |
| | Assaf_MB | Alval_MB | -0.315 | 0.159 | -1.983 | 0.426 |
| | Alval_MB | OBI_MB | 0.428 | 0.150 | 2.858 | 0.043* |
| BB vs MB | OBI_BB | OBI_MB | 0.062 | 0.096 | 0.645 | 1.000 |
| | Assaf_BB | Assaf_MB | -0.062 | 0.094 | -0.654 | 1.000 |
| | Alval_BB | Alval_MB | -0.424 | 0.095 | -4.469 | 9.42e-05*** |
| | OBI_BB | Alval_MB | -0.253 | 0.157 | -1.606 | 0.867 |

**Table S6.** Results of the multiple comparisons (with Holm correction) between hazard ratios obtained for different *T. evansi* population (BR, GH, and QL) among the different treatments (BB: *Beauveria bassiana*; MB: *Metarhizium brunneum*; C: control). Hazard ratios of infection by each fungus is estimated relative to the control within each population.

| | Factor 1 | Factor 2 | Estimate | Std. Error | z value | Pr(>\|z\|) |
|---|---|---|---|---|---|---|
| Controls | BR_C | GH_C | -0.263 | 0.218 | -1.205 | 0.914 |
| | GH_C | QL_C | -1.370 | 0.189 | -7.236 | 3.70e-12*** |
| | QL_C | BR_C | 1.633 | 0.201 | 8.145 | 4.44e-15*** |
| BB | BR_BB | GH_BB | 0.207 | 0.207 | -0.254 | 0.914 |
| | BR_BB | QL_BB | 1.435 | 0.188 | 7.631 | 2.10e-13*** |
| MB | BR_MB | GH_MB | -0.289 | 0.210 | -1.375 | 0.846 |
| | BR_MB | QL_MB | 1.176 | 0.192 | 6.117 | 6.68e-9*** |
| BB vs MB | BR_BB | BR_MB | 0.161 | 0.104 | 1.556 | 0.718 |
| | GH_BB | GH_MB | -0.075 | 0.097 | -0.775 | 0.914 |
| | QL_BB | QL_MB | -0.098 | 0.094 | -1.039 | 0.914 |

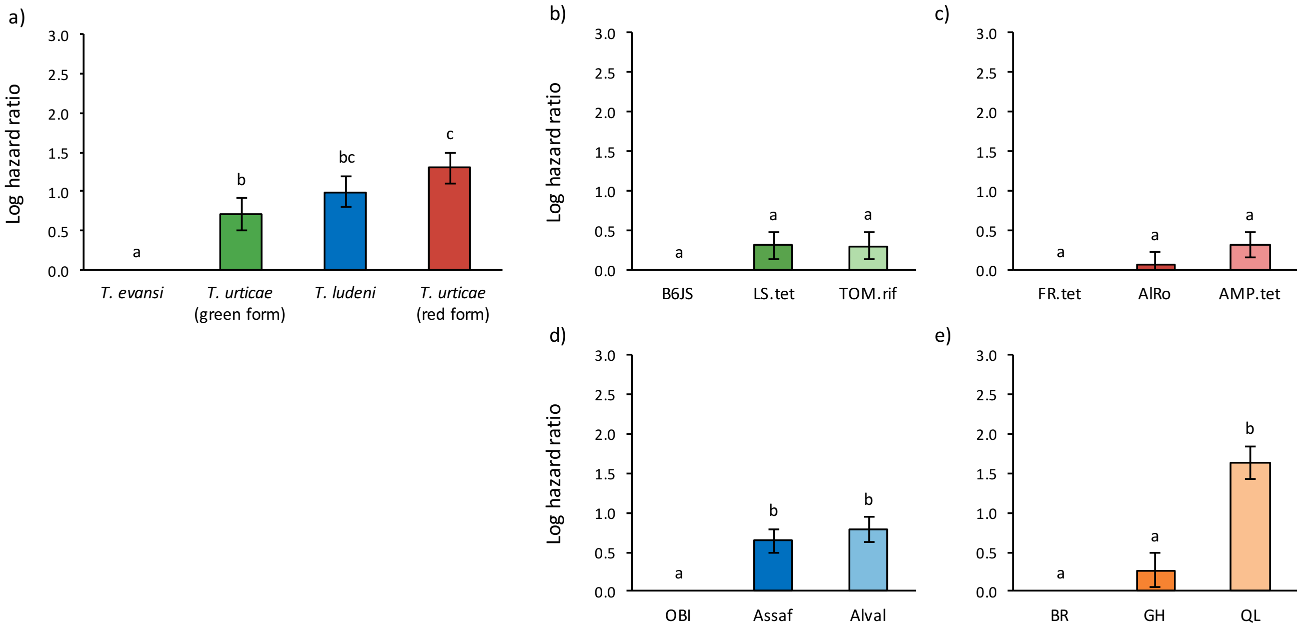


**Figure S1** Relative mortality (log hazard ratio ± s.e.) of the controls (sprayed with Tween 20 only) of (a) each species, (b) each population of the green form of *T. urticae*, (c) each population of the red form of *T. urticae*, (d) each *T. ludeni* population, and (e) each *T. evansi* population. Identical letters indicate non-significant differences between treatments at the 5% level (multiple comparisons with Holm correction).
